# Differential attention-dependent adjustment of frequency, power and phase in primary sensory and frontoparietal areas

**DOI:** 10.1101/697615

**Authors:** Nina Suess, Thomas Hartmann, Nathan Weisz

## Abstract

Continuously prioritizing behaviourally relevant information from the environment for improved stimulus processing is a crucial function of attention. Low-frequency phase alignment of neural activity in primary sensory areas, with respect to attended/ignored features has been suggested to support top-down prioritization. Phase adjustment in frontoparietal regions has not been widely studied, despite general implication of these in top-down selection of information. In the current MEG study, we investigated how ongoing oscillatory activity of both sensory and non-sensory brain regions are differentially impacted by attentional focus. Participants performed an established intermodal selective attention task, where low-frequency auditory (1.6 Hz) and visual (1.8 Hz) stimuli were presented simultaneously. We instructed participants to either attend to the auditory or to the visual stimuli and to detect targets while ignoring the other stimulus stream. As expected, the strongest phase adjustment was observed in primary sensory regions for auditory and for visual stimulation, independent of attentional focus. We found greater differences in phase locking between attended and ignored stimulation for the visual modality. Interestingly, auditory temporal regions show small but significant attention-dependent entrainment even for visual stimulation. Extending findings from invasive recordings in non-human primates, we demonstrate an effect of attentional focus on the phase of the entrained oscillations in auditory and visual cortex which may be driven by phase locked increases of induced power. In contrast to the effects in sensory areas, attentional focus adjusted the peak frequencies in nonsensory areas. Spatially these areas show a striking overlap with core regions of the dorsal attention network and the frontoparietal network. This suggests that these areas prioritize the attended modality by optimally exploiting the temporal structure of stimulation. Overall, our study complements and extends previous work by showing a differential effect of attentional focus on entrained oscillations in primary sensory areas and frontoparietal areas.

## 1. Introduction

Optimal processing of sensory stimuli from the environment is a crucial prerequisite for goal-directed adaptive behaviour. Interestingly, most behaviourally relevant sensory events like speech, show quasi-rhythmicity at slow frequencies (Ghitza, 2011), enabling the brain to track those slow modulations. Alongside attention-dependent amplitude modulation in early sensory areas (Alho et al., 1992; Bidet-Caulet et al., 2007; Mehta et al., 2000), a complementary process seems to exploit the temporally predictable structure of the input (Lakatos et al., 2007). In this context, neural entrainment (or phase alignment) of slow oscillations has been proposed to be an important process (Lakatos et al., 2008) for optimally adjusting cycles of neural excitability to the attended or ignored input. This process of adjusting phase “in a broad sense” ((Obleser & Kayser, 2019); this is what we sometimes refer to “entrainment” in this manuscript for the sake of simplicity) in sensory areas seems to be top-down mediated, putatively by higher-order brain regions which underlie sensory selection. Indeed several studies propose an interaction between primary sensory and non-sensory ‘control’ regions (Gazzaley & Nobre, 2012). In particular, areas of the so-called dorsal attention network (DAN), which involves connections between the intraparietal sulcus (IPS) and frontal eye fields have been implicated in top-down mediated target selection and target detection of bottom-up distinctiveness between stimuli (Buschman & Miller, 2007; Corbetta & Shulman, 2002). Additionally, areas in the frontoparietal network also act upon attentional modulation, with the IPS being involved in processing surprise targets, while the anterior cingulate cortex and the dorsolateral prefrontal cortex interact when guiding attention (Wang et al., 2009). Currently the effects of attention on entrained oscillations in higher-order (non-sensory) regions are unknown. To fill this gap, this study investigated entrained oscillatory activity using MEG. We used an established intermodal selective attention task which has been widely used (Calderone et al., 2014) to study attentional effects on entrained oscillations in primary sensory areas of the brain. This has previously been demonstrated using invasive recordings in nonhuman primates (Lakatos et al., 2016), as well as in patients with epilepsy (Gomez-Ramirez et al., 2011). Individuals were simultaneously presented with an auditory and a visual stimulus stream, and instructed to attend to one of these, while ignoring the other. To track the influence of attention on entrained oscillations separately for both modalities, each modality was stimulated with a different frequency. We replicate established attentional effects on phase alignment in primary sensory areas (e.g. Lakatos et al., 2016) and additionally find strong attention-dependent phase effects on the envelope of broadband induced power. Furthermore, properties of attentional phase effects in areas underlying flexible switching between modalities were investigated. Regions, strikingly overlapping with core regions of the dorsal attention system (Corbetta et al., 2008; Shulman et al., 2010; Szczepanski et al., 2013) and frontoparietal regions (Ptak, 2012) adjusted their stimulus-driven peak frequency flexibly to the rhythm of the attended modality. This finding is an important advancement in integrating reports of attentional effects on entrained oscillations in primary sensory areas with high-level and putatively supramodal processes of the dorsal attention network and the frontoparietal network.

## 2. Materials and Methods

### 2.1 Participants

We recruited 33 participants (15 females; 4 left-handed; mean age: 26.3 years; SD: 7.9 years) for the experiment. Two subjects had to be excluded, the first one because there were problems with the head digitization and the second participant was not able to perform the visual task. All participants had normal or corrected-to-normal eyesight, normal hearing and no neurological disorders. All participants received either a reimbursement of €15 for their time, or if they were Psychology students, they received credits for their participation. All participants signed an informed consent form. The experimental procedure was approved by the Ethics Committee of the University of Salzburg.

### 2.2 Stimuli

Participants were presented with an auditory and a visual stimulus stream simultaneously in each block. Before the main experiment, a 4-minute training session was carried out for auditory and visual targets separately to determine the perception threshold at which 75% of the target stimuli were detected by the participants. This was achieved using a Bayesian active sampling protocol to estimate the model parameters of the psychometric function (Kontsevich & Tyler, 1999; Sanchez et al., 2016). The procedure was implemented using the VBA toolbox in Matlab (Daunizeau et al., 2014). The procedure was carried out in the same environment and with the same hardware as the final experiment. The visual standard stimuli were black circles with a visual angle of 3.5° on a grey screen. The visual targets were different from the standard stimuli in terms of colour, meaning that according to the adjusted perception threshold a grey circle instead of a black circle was presented. The threshold for the deviant was adjusted between RGB values from 0 to 96. Visual stimuli were back-projected for 25 ms on a translucent screen in the centre of the screen by a Propixx DLP projector (VPixx technologies, Canada) with a refresh rate of 120 Hz per second and a screen resolution of 1920 x 1080 pixels. The translucent screen was placed ~110 cm in front of the participant and had a size of 74 cm. Auditory standard stimuli were 440 Hz pure tones of 25 ms duration that were presented binaurally with MEG-compatible in-ear headphones (SOUNDPixx, VPixx technologies, Canada). The auditory targets were different from the standard tones in frequency, meaning that the targets were higher. The threshold for the deviant was adjusted between 440 and 550 Hz. The presented frequency for the targets was also determined in the aforementioned staircase.

### 2.3 Procedure

In the main experiment, participants performed 10 blocks of a selective intermodal attention task (Figure 1). Participants were instructed before each block to attend to either the auditory stream and detect the deviant tone which was higher while ignoring the simultaneously presented visual stimuli (“attend auditory”) or alternatively, to attend to the visual stream and detect the deviant circle which was brighter while ignoring the presented auditory stream (“attend visual”). The “attend visual” and “attend auditory” blocks were alternated (see also Besle et al., 2011). The different stimulus streams were presented with differing SOAs to avoid having a constant temporal relationship between visual and auditory stimulus streams to allow for independent tagging of the frequency in the regions of interest (auditory and visual primary sensory areas, respectively) (see also Lakatos et al., 2016). The visual stream was programmed to have a 1.8 Hz repetition rate but since our projector was limited to a refresh rate of 120 Hz, our repetition rate resulted in 1.79 Hz with a SOA of 558.1 ms. For the purpose of simplification, we from now on refer to the visual stimulation rate as 1.8 Hz. The auditory stream had a SOA of 625 ms (1.6 Hz repetition rate). These frequencies were chosen to correspond to the delta frequency range (1-3 Hz) of ongoing brain oscillations to match the frequencies used by Lakatos et al. (2016) in a similar paradigm. The response time window matched the SOA between the stimuli (558.1 ms for the visual stream and 625 ms for the auditory stream, respectively). The responses were given with MEG-compatible response boxes (ResponsePixx, VPixx technologies, Canada). If the person took longer for a response, the trial was classified as a miss. All participants were instructed to use their left thumb for responding. Each run was 4 minutes long, resulting in 384 auditory stimuli and 432 visual stimuli. Out of those stimuli, 10% were targets (38 and 43 for every block, respectively, resulting in 215 visual targets and 190 auditory targets for every subject). The whole experiment lasted about 1.5 hours including preparation and staircase procedure. The experimental procedure was programmed in Matlab with the Psychtoolbox 3 (Brainard, 1997) and an additional class-based abstraction layer (https://gitlab.com/thht/th_ptb) programmed on top of the Psychtoolbox.

**Figure 1:**
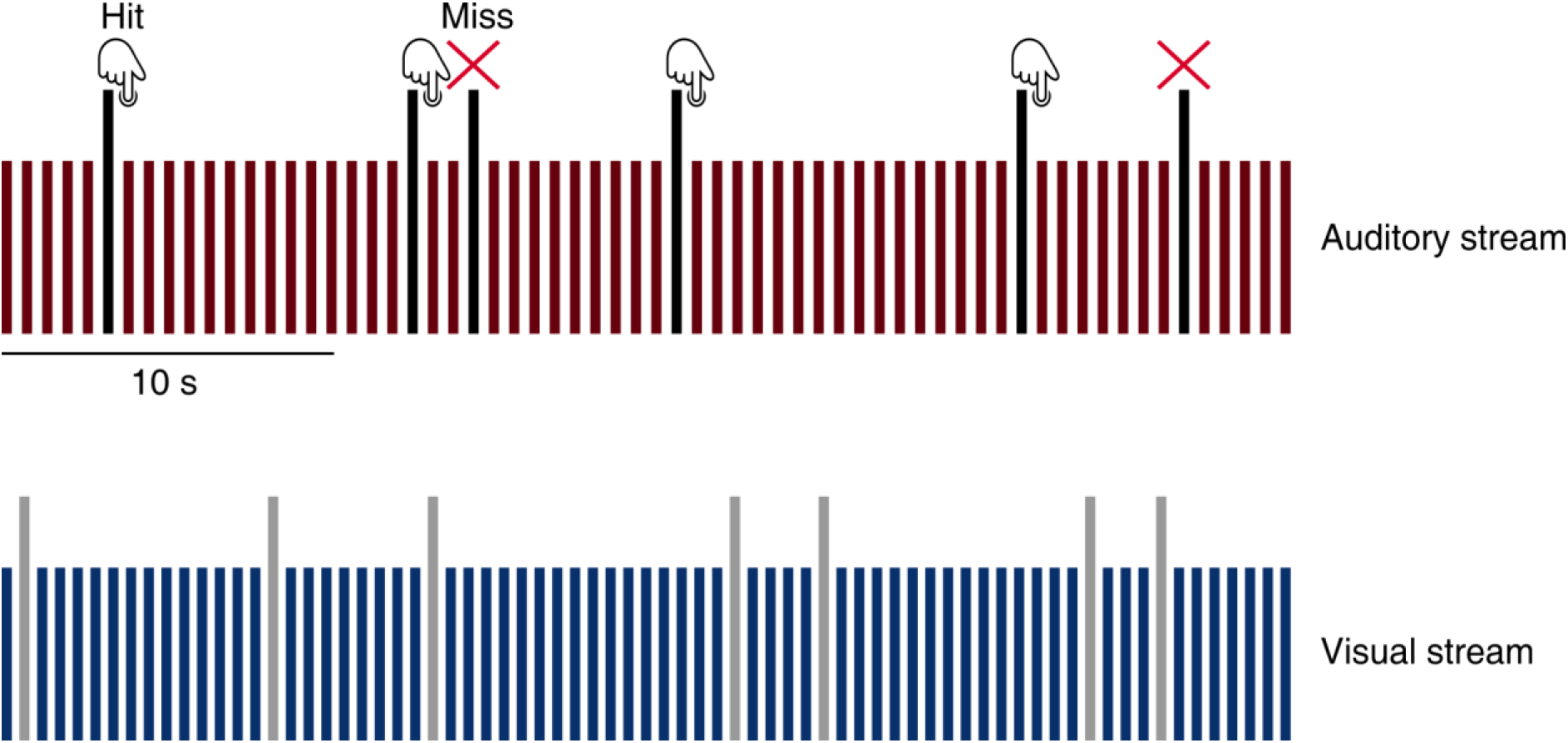
Intermodal selective attention task. For the auditory stream (1.6 Hz repetition rate), red bars represent standard tones, black bars represent target tones. For the visual stream (1.8 Hz repetition rate), blue bars represent standard stimuli, gray bars represent target stimuli. Performance was recorded by pushing a button right after target appearance. For visualization purposes we just depicted the “attend auditory” condition. Since false alarms did not occur during the experiment, they are not shown in this figure.

### 2.4 Data acquisition

Brain activity was measured using a 306-channel whole head MEG system (Neuromag TRIUX, Elekta) with a sampling rate of 1000 Hz. This system uses 204 planar gradiometers and 102 magnetometers. Before entering the magnetically shielded room (AK3B, Vakuumschmelze, Hanau, Germany), the head shape of each participant was acquired using about 300 digitized points on the scalp was acquired, including fiducials (nasion, left and right pre-auricular points) with a Polhemus Fastrak system (Polhemus, Vermont, USA). After acquisition, the continuous MEG data was preprocessed off-line with the signal space separation method from the Maxfilter software (Elekta Oy, Helsinki, Finland) to correct for different head positions across blocks and to suppress external interference (Taulu et al., 2005). The head position of each individual subject relative to the MEG sensors was controlled once before each experimental block. Additionally, vertical and horizontal eye movement and electrocardiographic data were recorded and used for artefact detection.

### 2.5 Data analysis

#### 2.5.1 Preprocessing

Acquired datasets were analysed using the Fieldtrip toolbox (Oostenveld et al., 2011). The maxfiltered MEG data was highpass-filtered at 0.1 Hz using a finite impulse response (FIR) filter (Kaiser window, order 36222). For extracting physiological artefacts from the data, 60 principal components were calculated from the high-pass filtered data at 0.1 Hz. Via visual inspection, the components showing eye-movements, heartbeat and external power noise from the train (16.67 Hz) were removed from the data. We removed on average 4 components per subject (*SD* = 1). To be able to extract the Fourier coefficients for the exact frequency of interest (1.6 Hz for auditory stimulation and 1.79 Hz for visual stimulation), we chose a window length of five cycles per frequency of interest (cpf) as this yields the necessary spectral resolution at low frequencies. We thus extracted 3.125 seconds for each auditory trial and 2.79 seconds for each visual trial, data centered at stimulus onset. The extracted data was then multiplied by a hanning taper to reduce spectral leakage. Finally, we applied a Fourier Transform to each of the tapered single trials to obtain the respective complex fourier coefficients.

#### 2.5.2 Source projection of MEG data

We used a standard structural brain from the Montreal Neurological Institute (MNI, Montreal, Canada) and warped it into the individual head shape (Polhemus points) to match the individuals fiducials and head shape reference landmarks as accurately as possible. A 3-D grid with 1 cm resolution and 2982 voxels based on an MNI template brain was morphed into the brain volume of each participant. This allows group-level averaging and statistical analysis as all the grid points in the warped grid belong to the same brain region across subjects. These aligned brain volumes were also used for computing single-shell head models and leadfields. By using the leadfields and the common covariance matrix (pooling data from all blocks), a common LCMV (Veen et al., 1997) beamformer spatial filter was computed. We then applied the spatial filter to the complex fourier coefficients obtained in the previous step to find the estimated complex source signal (Bardouille & Ross, 2008). The further analysis was limited to the frequency band of interest of 1-3 Hz.

#### 2.5.3 ITC analysis

To characterize the magnitude of entrainment across trials, we calculated the intertrial coherence (ITC) at the respective frequencies of interest for all trials, including hits and misses (1.6 Hz for “attend auditory” and 1.8 Hz for “attend visual” condition). We therefore extracted the Fourier coefficient at every voxel and calculated the average of the lengths of the normalized single-trial vectors, which then results in a single resultant vector. The length of the resultant vector can reach a number between 0 and 1 and represents the similarity of phases across trials. Higher values indicate that the phase distribution of the trials at a given time-point is clustered more closely around the angle of the mean resultant vector, while lower values indicate that the phase distribution of the trials at that given-time point are not clustered around the mean resultant vector.

#### 2.5.4 Phase differences

For calculating phase differences between attended and unattended stimulus streams we used the phase opposition sum (POS) introduced by VanRullen (2016), which is a measure for the consistency of phase differences over trials. Phase opposition is defined as the difference in angles between two waves that are oscillating with the same temporal resolution. Maximal phase opposition is reached when at one particular time point the waves show a 180° phase difference. The POS is calculated by using the following formula:

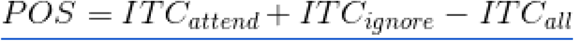

ITC_attend_ stands for the attended condition (either auditory or visual stimulation) and the ITC_ignore_ stands for the ignored condition (attend auditory – visual stimulation or attend visual – auditory stimulation). ITC_all_ takes into account all the ITC values calculated in the attended and not attended stimulus stream. POS can be similarly interpreted as the PBI (phase bifurcation index) (Busch et al., 2009), meaning that the value will be positive when the ITC of each group is higher than the overall ITC value.

### 2.6 Statistical analysis

#### 2.6.1. Region-of-interest analysis

We defined our functional regions-of-interest (ROI) by extracting the voxels that reached at least 75% of the maximum ITC value in the “attend auditory” condition and in the “attend visual” condition. The resulting areas corresponded anatomically to the temporal and occipital cortices, respectively (Figure 2a). We then averaged averaged the chosen voxels for every subject. For clarifying the relationship between conditions (factors: “attend” and “ignore”), region of interest (factors: “visual” and “auditory”) and stimulation (factors: “visual” and “auditory”), we performed a 2 x 2 x 2 repeated-measures ANOVA with the following factors; CONDITION x ROI x STIMULATION.

**Figure 2:**
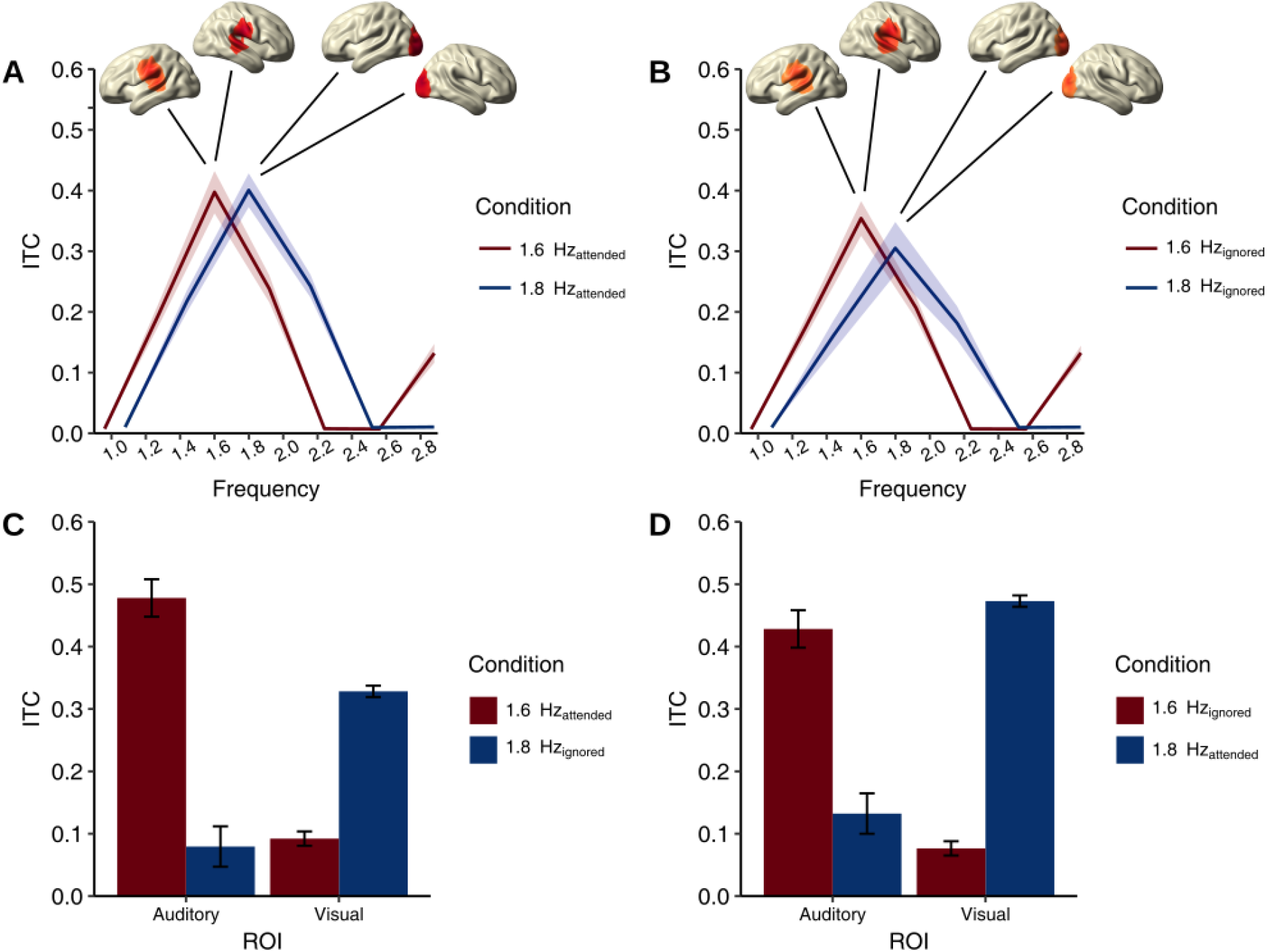
Entrainment effect in primary sensory areas. A) Comparison of the means of the attended stimulation (red line = auditory and blue line = visual) for voxels reaching at least a threshold of 75% of the maximum ITC value extracted from temporal areas (for auditory stimulation, 1.6 Hz) and from occipital areas (for visual stimulation, 1.8 Hz). Both areas show similar ITC when attending to the preferred stimulation. B) Comparison of the means of the ignored stimulation (red line = auditory, blue line = visual) for voxels reaching at least a threshold of 75% of the maximum ITC value extracted from temporal areas (for auditory stimulation, 1.6 Hz) and from occipital areas (for visual stimulation, 1.8 Hz). Both areas show a tendency to also track ignored stimulation, but less pronounced for the visual stimulation. C) Differences between auditory (1.6 Hz, red bars) and visual (1.8 Hz, blue bars) stimulation depicted for the voxel with the highest ITC value in the “attend auditory” condition extracted from temporal areas for the auditory ROI and from occipital areas for the visual ROI. High entrainment in temporal areas when attending to auditory stimulation (1.6 Hz), but also high entrainment in occipital areas when ignoring the simultaneously presented visual stimulation (1.8 Hz). D) Differences between auditory (1.6 Hz, red bars) and visual (1.8 Hz, blue bars) stimulation depicted for the voxel with the highest ITC value in the “attend visual” condition extracted from occipital areas for the auditory ROI and from auditory areas for the visual ROI. High entrainment in occipital areas when attending to visual stimulation (1.8 Hz, blue), but also high entrainment in temporal areas when ignoring the simultaneously presented auditory stimulation (1.6 Hz, red). Error bars represent 1 SEM.

#### 2.6.2 Phase opposition sum (POS)

For our frequency of interest (1.6 Hz for the auditory stimulation and 1.8 Hz for the visual stimulation) we individually computed an ITC value for every voxel. We used a 1 cm grid with 2982 voxels and then used the proposed POS analysis by VanRullen (2016) to compute the values for each voxel separately. We used a permutation test containing 1000 permutations where every trial was randomly assigned to the ITC attend or the ITC ignore condition and after every permutation, the POS was recomputed for every voxel. The final p-value shows the proportion of permutations with a higher measure than in the original data. After calculating the p-values on a single-subject basis, we then combined the p-values for every voxel using the Fisher’s method where the p-values are combined in the log domain and it is assumed that the null hypothesis follows a chi-square distribution. We then applied a Bonferroni correction on a 5% level for the combined p-values.

#### 2.6.3 Modality-independent attention effect

We were interested which regions adjust the frequency of their entrained oscillations relative to the attentional focus. For this purpose, we first calculated the mean ITC over the attended streams (in the “attend auditory” condition the values calculated for the auditory trigger (1.6 Hz) and in the “attend visual” condition the values calculated for the visual trigger (1.8 Hz)). We continued with the same procedure for the ignored stimulation (taking the mean ITC values from the respective ignored stimulus streams). The following formulas summarizes the computations:

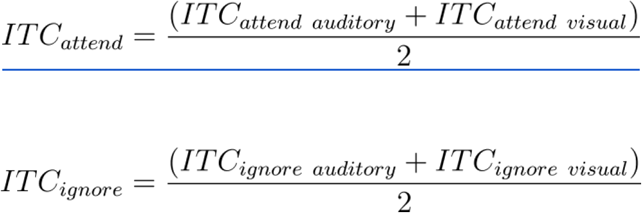

To contrast the ITC values from the attended stimulation with the ITC values from the ignored stimulation, we compared all voxels using a dependent-samples t-test with Bonferroni correction to control for multiple comparisons on a 5% level.

## 3. Results

### 3.1 ITC values are highest in primary sensory areas for modality specific (auditory or visual) stimulation

We first performed a descriptive whole brain analysis to investigate which parts of the brain show the highest entrainment to the stimulation. We found the highest entrainment for the stimulation frequency in temporal areas (1.6 Hz, ITC_attend_ = 0.478, MNI coordinates [70.0 – 20.0 20.0]) and occipital areas (1.8 Hz, ITC_attend_ = 0.473, MNI coordinates [−20.0 −100.0 0.0]) (Figure 2a). These areas of visual and auditory cortex remain actively entrained while input from the matched sensory modality is ignored, albeit at a somewhat lower level (ITC_ignore_ = 0.428 in temporal areas (Figure 2c) and ITC_ignore_ = 0.328 in occipital areas (Figure 2d)), indicating that the temporal features of the input are still faithfully tracked, even when not attended to. As expected, there was low entrainment to visual stimulation in the auditory cortex (ITC_ignore_ = 0.079 (Figure 2c), ITC_attend_ = 0.132 (Figure 2d)) and the same low entrainment pattern in the visual cortex for auditory stimulation (ITC_attend_ = 0.092 (Figure 2c), ITC_ignore_ = 0.077). Descriptive inspection revealed slightly higher entrainment for the attended not-matching modality, especially in auditory areas, where attending to visual stimulation also lead to higher entrainment in auditory cortex. For descriptive purposes, all values in Figures 2c and 2d described for sensory modalities are from the voxel with the highest value in the “attend” condition.

To statistically assess differences depicted in Figure 2, we performed a repeated-measures ANOVA with the factors CONDITION x ROI x STIMULATION. We extracted the voxels depicted in Figure 2a (for both auditory and visual stimulation) and averaged them for statistical analyses. We found a significant difference between conditions (“attend” vs. “ignore”: F_(1,30)_ = 35.19, *p = 1.04e-08,* Figure 3a) and a significant difference between auditory and visual ROIs (F_(1,30)_ = 4.52, *p = 0.035,* Figure 3b). We also found a clear interaction effect for stimulation and ROI, (F_(1,30)_ = 652.05, *p < 2e-16*), showing that there is high entrainment in auditory cortex for auditory stimulation and high entrainment in visual cortex for visual stimulation (Figure 3c). Furthermore, we found a significant interaction effect between condition and stimulation (F_(1,30)_ = 7.45, *p = 0.007)* depicted in Figure 3d, which shows after further analysis (ANOVA with the factors CONDITION x STIMULATION) that while for the auditory stimulation there was smaller, but still significant difference between the attended and ignored condition (F_(1,30)_ = 4.48, *p = 0.036),* there was a big and significant difference for the visual stimulation depending on the condition (F_(1,30)_ = 43.91, *p = 1.03e-09*). This shows that the brain entrains differentially to the attended or ignored stimulation, but this distinction is more prominent in the visual domain. This also suggests that the auditory cortex shows a tendency to temporally align activity to visual information when attended (e.g. Besle et al., 2011). Overall, our results show that sensory cortices entrain to rhythmic sensory input regardless of whether the input is attended or not, but there is increased entrainment of the attended stimulation. Interestingly, we find higher differences between the “attend visual” and the “ignore visual” condition independent of cortical regions, suggesting that the auditory cortex also shows modulation at the visual frequency. To explore this further, we investigated phase differences for different modalities separately.

**Figure 3:**
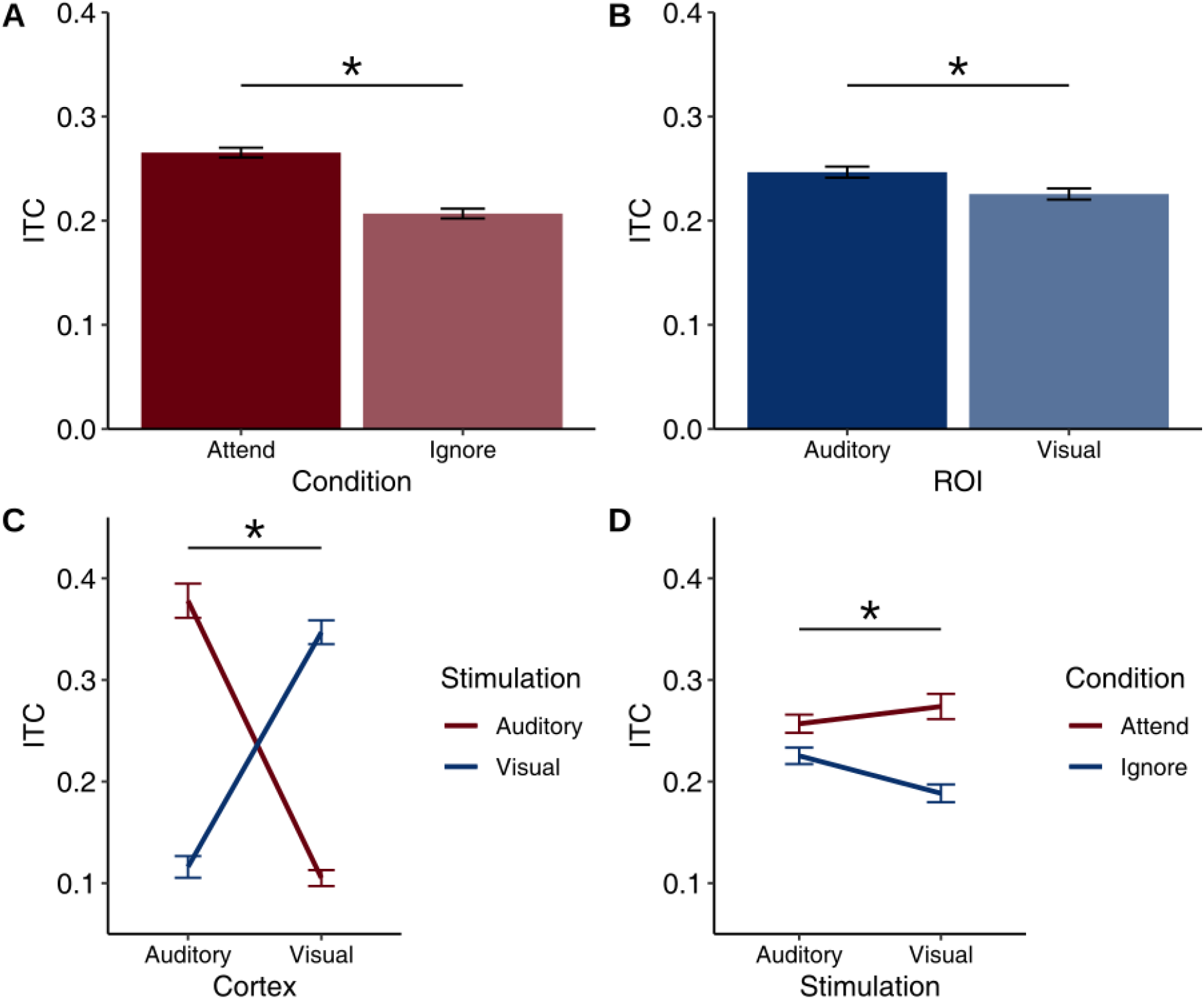
Main effects and interactions of the repeated measures ANOVA. A) Significant main effect for attended vs. ignored stimulation. Higher ITC values when participants are instructed to attend to a certain stimulation. B) Higher ITC values in the auditory ROI in general regardless of stimulation or condition. C) Significant interaction between stimulation (auditory and visual) and ROI. High ITC values in auditory ROI and low ITC values in visual ROI when presenting auditory stimulation, while having high ITC values in visual ROI and low ITC values in auditory ROI when presenting visual stimulation. D) Significant interaction between conditions (“attend” and “ignore”) and stimulation. Low differences in the conditions when presenting auditory stimulation, but high differences in conditions when presenting visual stimulation. Error bars represent 1 SEM for within-subject designs (Morey, 2008).

### 3.2 Attentional influences on phase in primary sensory areas

After finding entrainment effects most prominent in sensory areas, we were interested in how attention shapes the phase of stimulus-driven slow oscillations in the brain. It has been previously established in primate studies that shifting the attentional focus also results in prominent phase shifts between the attended and ignored stimulation (e.g. Lakatos et al., (2016), Lakatos et al., (2013)), but for non-invasively recorded data there hasn’t been well established evidence yet, especially in the visual domain. Using the fine temporal resolution of MEG, oscillatory responses to attended and ignored stimulus streams were extracted and compared. For a group-level analysis, we calculated the phase opposition sum (POS) proposed by VanRullen (2016) on a single subject basis and then calculated the combined p-value over all subjects for each voxel separately. The POS contrasts the attended and ignored ITC values separately for auditory and visual stimulation. After Bonferroni correction, we still found significant widespread differences throughout the brain, so in order to describe the most prominent effects only the lowest 1% of the observed p-values are shown (separately for auditory and visual stimulation). This procedure revealed most consistent phase differences over primary sensory areas. These phase differences were most prominent in the left superior temporal sulcus (MNI coordinates [−60.0 −30.0 10.0], Figure 4a) for the auditory stimulation (*p = 8.5141e-50).* and most prominent for visual stimulation (*p = 5.073e-61)* in right occipital areas (MNI coordinates [10.0 −90.0 10.0], Figure 4c). Figure 4b and 4d depict the evoked response data bandpass-filtered between 1-2 Hz for the voxel with the lowest p-value for a single subject in auditory and visual cortex respectively. Descriptively these results are similar to monkey data, showing a clear phase difference for attended and ignored stimulation both in the auditory condition and in the visual condition when extracting the voxels with the lowest p-value. Interestingly, this depiction also reveals strong POS effects in sensorimotor cortex for the auditory stimulation, underlining the involvement of these regions in auditory rhythm processing (Chen et al., 2006).

**Figure 4:**
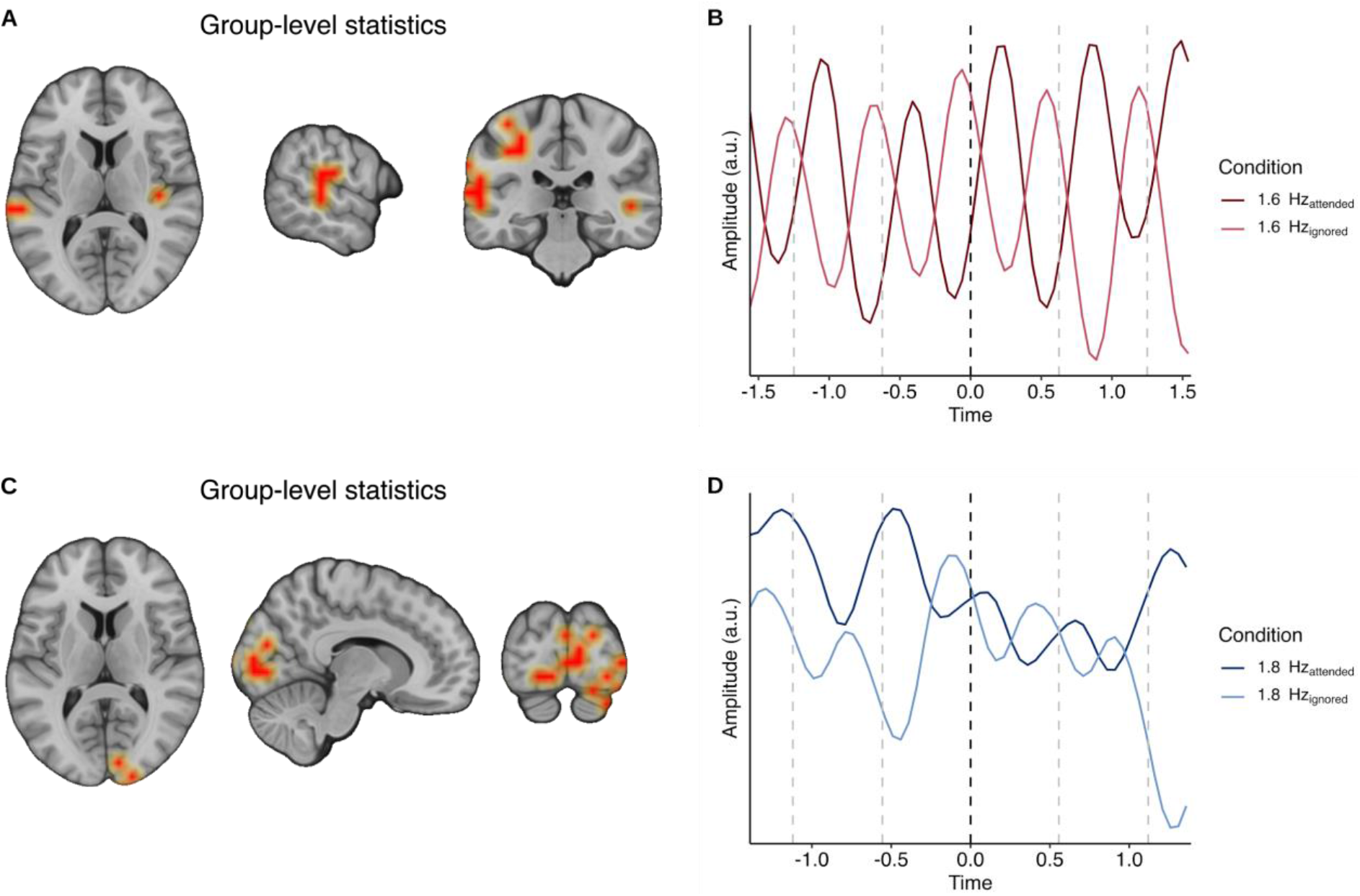
Visualization of phase opposition and source maps of p-values calculated for every voxel for respective stimulation frequency. A) Brain plots showing significant phase differences for the auditory stimulation present in voxels over left temporal areas and in the left superior temporal sulcus on a group level (MNI coordinates [−60.0 −30.0 10.0]). Marked (orange) voxels show the lowest 1% of significant p-values. B) Example single-subject bandpass-filtered timelocked data for the voxel with the lowest p-value showing phase differences during auditory stimulus presentation. C) Brain plots showing significant phase differences for visual stimulation in voxels over occipital areas on a group level (MNI coordinates [10.0 −90.0 10.0]). Marked (orange) voxels show the lowest 1% of significant p-values. D) Example single-subject timelocked bandpass-filtered data for the voxel with the lowest p-value showing phase differences during visual stimulus presentation.

This previously described results shows that in sensory regions attention adjusts the timing of neural activity, being captured in the POS measure. Since the stimulation related neural activity at a single-trial level could be linked to a “pure” phase modulation as well as power increases, our attention-related POS effect could in principle stem from both processes. In a follow-up analysis we decided to investigate to what extent single-trial (induced) power is entrained to stimulation frequencies and whether these modulations show analogous POS effects as described above. We calculated the single-trial power modulation for visual and auditory stimulation separately by first low-pass filtering the signal at 40 Hz and subsequently applying Hilbert-transformation in order to obtain the single-trial broadband power envelopes. These were subjected to the same POS analysis as described previously (from now on referred to as POS_Hilb_) for attended and ignored stimulation for the respective stimulation frequency (1.6 Hz for auditory stimulation and 1.8 Hz for the ignored stimulation) for the voxel showing the lowest p-value in our initial analysis (Figure 4a and c). We found a significant POS_Hilb_ between attended and ignored stimulation for auditory stimulation at the stimulation frequency (1.6 Hz, p = 2.489e-26) and for visual stimulation (1.8 Hz, p = 1.006e-40), showing that our “pure” phase modulation could also be due to latency shifts in power modulation at stimulation frequencies.

### 3.3 Fronto-parietal areas adjust entrained oscillation frequency in a supramodal manner

While sensory regions exhibited attentional modulations at frequencies used for entrainment, they were overall modality specific. The general role of the dorsal attention network in mediating top-down guided attention to stimulus features together with the fronto-parietal network being responsible for attentional control would imply a flexible supramodal process. Here we tested in a data-driven manner the existence of regions that adjust their slow oscillatory dynamics flexibly to the temporal and modality-specific structure of external input. We used the formula explained in 2.6.3. with which we calculated the mean over the attended and ignored ITC values independent from modalities and applied a dependent samples t-Test with Bonferroni correction over all voxels to compare the “attend” condition with the “ignore” condition. To illustrate that the entrainment effect for attention is corresponding to the dorsal attention network by Corbetta and colleagues (2008), we marked all the voxels showing an overlap with the dorsal attention network as reported in the parcellation approach by Gordon et al. (2016). Additionally, we investigated the activation in voxels overlapping with the frontoparietal network. These proposed areas involve in particular the frontal eye fields (FEF), the intraparietal sulcus (IPS), the anterior cingulate cortex (ACC) and the dorsolateral prefrontal cortex (DLPFC). Significant differences between the attended and ignored stimulation were found in a distributed set of regions, with maximum effects in parietal and frontal areas (highest *t*-value: *t*(30) = 9.538, p < 0.000, MNI coordinates [20.0 −50.0 60.0]). We find clear overlaps in intraparietal areas and the frontal eye field (Figure 6a and Figure 6b) which also show strongest t-values in our statistical analysis. Significant t-values overlap with 17.8% of the proposed dorsal attention network and with 15.4% of the frontoparietal network, while only 8.5% and 2% of the significant voxels overlap with the default network and the ventral attention network, respectively (Figure 5). This result shows that distinct areas entrain flexibly to the endogenously attended stimulation rate independent of the sensory modality.

**Figure 5:**
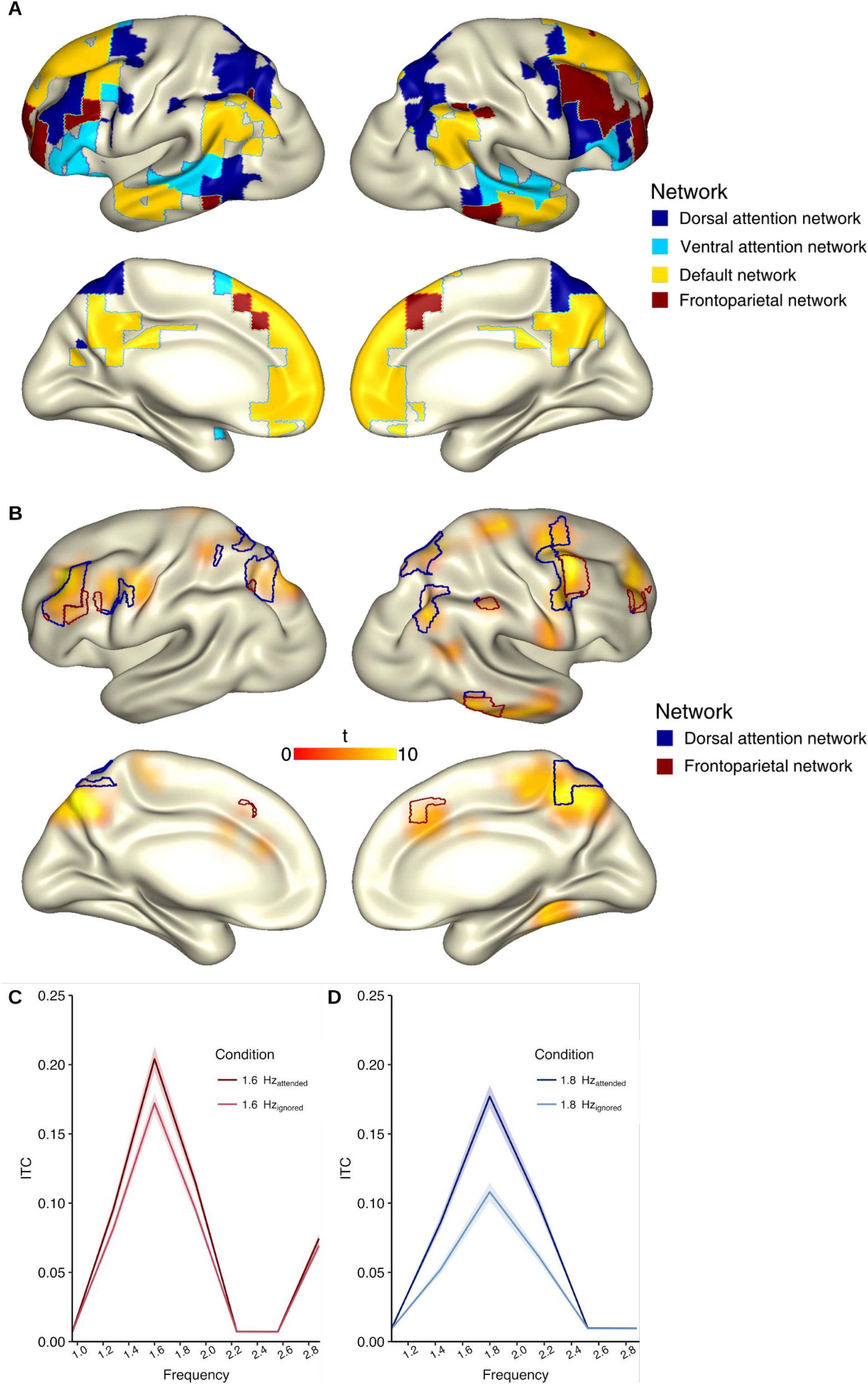
Main attention effect across conditions and comparison between brain areas. A) Different networks from the atlas proposed by Gordon et al. (2016). Shown are the dorsal attention network, the ventral attention network, the default network and the frontoparietal network. B) Comparison of attended and ignored stimulation independent from sensory modality (dependent samples t-Test, Bonferroni corrected). Highest value over right parietal areas, with expansion to frontal areas. Blue lines indicate overlap with the dorsal attention network, red lines indicate overlap with the frontoparietal network. C) Comparison of the means of the auditory stimulation (1.6 Hz, “attend auditory” vs. “ignore auditory”) for voxels extracted from the frontoparietal and the dorsal attention network as proposed in the aforementioned atlas. Higher ITC values for attended stimulation than for the ignored stimulation. D) Comparison of the means of the visual stimulation (1.8 Hz, “attend visual” vs. “ignore visual”) for voxels extracted from the frontoparietal and the dorsal attention network as proposed in the aforementioned atlas. Higher ITC values for attended stimulation than for the ignored stimulation irrespective of modality. Shaded error bars represent 1 SEM.

Plotting an average over all voxels corresponding to the frontoparietal network and the dorsal attention network for the stimulation frequency as well as for the neighbouring frequencies (between 1 and 3 Hz) revealed in total lower entrainment values compared to entrainment in sensory cortices for the corresponding modality (see Figure 2a and b). Still, voxels in the dorsal attention network and the frontoparietal network show significantly higher entrainment for the attended stimulation (Figure 5c for auditory stimulation and Figure 5d for visual stimulation), thus switching the peak frequency dependent on attentional focus. This higher entrainment is not specific for one modality, providing evidence for a modality-independent processing of sensory input in high-order areas.

## 4. Discussion

The current results provide insights into the differential impact of endogenous attention on entrained (or phase locked) oscillations of sensory regions and areas of the dorsal attention network (Corbetta & Shulman, 2002) and the frontoparietal network. As expected, phase adjustment is strongest in visual and auditory areas for the respective stimulation modality, independent of attention. In accordance with the primate literature (Lakatos et al., 2013, 2016; O’Connell et al., 2014) attending to or ignoring a stimulus leads to a phasic difference in primary sensory areas for the specific input modality, which could also be due to latency shifts in power modulation. Despite the small ITC effect in auditory cortex at the visual stimulation rate, the attentional effects in primary sensory areas were largely modality specific. Going beyond previous reports, we show that core areas of the dorsal attention network and the frontoparietal network flexibly adjust their entrained frequency to the attended stimulation independent from sensory input. This flexible adaptation is likely crucial in optimizing sensory processing of the selected input.

### 4.1 Entrainment is amplified by attention in primary sensory areas in a largely modality-specific manner

Calculating the grand average over all subjects and all voxels revealed as expected highest phase adjustment in primary sensory areas (Figure 2). We also found the highest modulation of entrainment for the respective stimulation frequency (1.6 Hz for auditory stimulation over temporal areas and 1.8 Hz for visual stimulation over occipital areas), which is in line with previous intersensory attention tasks conducted with both human individuals and with primates (Cravo et al., 2013; Gomez-Ramirez et al., 2011; Lakatos et al., 2008, 2009). Even with day to day multi-sensory input such as that from movies, sensory areas are able to entrain to the low-frequency properties of the stimulation. Interestingly, the auditory cortex shows an increase at the visual stimulation rate when visual input is attended to, indicating that it also tracks temporal aspects of the visual input. This result is in line with other studies showing an effect of visual input on auditory processing (Besle et al., 2011; Lakatos et al., 2016; Luo et al., 2010), proposing a crossmodal system for integrating different properties of the signal (Ghazanfar et al., 2005). Another feature that has been shown to be specific for the auditory cortex is the processing of rhythmicity (Zatorre et al., 2007). Synchronization of internal oscillatory properties to external stimuli in both the auditory cortex and motor cortex is important, especially in complex musical performances, (Chen et al., 2006), suggesting that both the auditory and motor cortex process the rhythmicity of environmental cues independent from modality to sample sensory input for optimal perception. However, in light of the results described for frontoparietal regions, it should be emphasized that despite the attentional increase of ITC at the visual stimulation rate in auditory cortex, the values are still only a fraction of the ITC to the auditory stimulation rate and this is so even when only considering the ignored condition (ITC_ignore_ = 0.428 for auditory stimulation vs. ITC_attend_ = 0.132 for visual stimulation in auditory cortex). Thus, the overall strength of entrainment (assessed via ITC) in sensory regions shows a clear preference for the corresponding input modality and this response is amplified by attention.

### 4.2 Attention adjusts low-frequency phase and power in primary sensory areas

Further to an amplification of sensory responses, and given the predictable rhythmicity of the stimulation, selective attention can also exploit temporal information to optimally align excitable phases according to the occurence of the prioritized or ignored features. This has previously been shown in invasive recordings to lead to marked phase alignment of entrained frequencies in primary sensory regions (Lakatos et al., 2008, 2016). These phasic modulations provide an optimal processing window and differentiate between perception and non-perception (Lakatos et al., 2007, 2009; Monto et al., 2008; Schroeder et al., 2010) and can also lead to faster reaction times (Stefanics et al., 2010). Calculating the POS for attended and ignored stimulation on a group-level basis revealed significant phase differences which were most prominent over corresponding sensory cortices on a group-level. The findings are in line with literature on recordings from monkey cortices (Lakatos et al., 2009, 2016; O’Connell et al., 2014) where counter-phase entrainment depending on attention has been reported. Shifts of entrained oscillatory rhythms by attention has been reported in occipital areas in nonhuman primates (Lakatos et al., 2008) further supporting the same mechanism for attentional selection in visual and auditory modalities (for invasive recordings in humans see Besle et al. (2011)). However, distinguishing “pure” phase modulation (to which entrainment refers to in a “narrow sense”; see Obleser & Kayser (2019)) is difficult when time – locked power modulations go along in parallel. We followed up this issue by Hilbert transforming the single trials before again calculating the POS using the power envelope. We found systematic shifts in these phase locked power modulations between the attended and ignored stimulation. This result does not exclude the possibility that there are still “pure” phase differences as both power and phase modulations contribute to evoked potentials (Fuentemilla et al., 2008). Until now, studies looking at low-frequency phase modulations in primary sensory areas mostly focused on one modality-specific region and mainly on phasic modulations in auditory cortex using invasive recordings (Lakatos et al., 2008). Our study non-invasively captures the attentional phase adjustment of entrained oscillations simultaneously in auditory and visual cortex, underlining previous reports that this mechanism is a versatile process across sensory modalities when prioritizing attended information. Going beyond the previous invasive studies, our work also points to attentional phase adjustment effects also in sensorimotor cortex with respect to the auditory stimulation. This effect is compatible with models that hold an important role of motor-cortical areas in processing of auditory rhythm information (Zatorre et al., 2007). To what extent this motor effect is functionally relevant in the context of this task needs to be investigated. In both primary sensory and sensorimotor cortices, the question arises whether “pure” phase adjustment is exclusively responsible for attention shifting or if power modulation and phase alignment contribute equally to the differentiation between attended and ignored stimulation.

### 4.3 The dorsal attention system flexibly adjusts its entrained frequency to the selected modality

Our previous results show that with regards to the magnitude and phase of entrained oscillations, attentional modulations in sensory regions are largely modality specific. Flexible endogenous selection of attended features across different sensory modalities would however require supramodal processes. Here we show for the first time that core regions of the dorsal attention network (DAN) and the frontoparietal network differentially entrain to attended temporal properties of sensory stimulation. While the right intraparietal sulcus, (involved in orientation to person, place, and time), registers salient events in the environment not only in the visual, but also in the auditory and tactile modalities (Downar et al., 2000; Lee et al., 2014), the inferior parietal lobe seems to be responsible for attention shifting, switching and the maintenance of attention (Ptak, 2012). Even when shifting attention voluntarily between visual or auditory input, the brain shows increased activation in posterior parietal and superior prefrontal cortices (Shomstein & Yantis, 2004), highlighting the importance of those areas for attentional control functions. These findings are supported and most importantly extended by our results (Figure 5) as we show entrainment to exogenous stimulation in parietal and frontal areas corresponding to the dorsal attention system (Corbetta et al., 2008) and the frontoparietal network (Marek & Dosenbach, 2018). Our results add to previous reports by showing that the regions of key attentional networks can flexibly switch between modalities and tune their dominant tracking speed to the to-be-attended frequency, further supporting optimal stimulus processing. This seems to be of particular importance as the delta frequency is the basis for modulating faster rhythms in the brain (Lakatos et al., 2005; Schroeder et al., 2008). We argue that slow-frequency modulations (at least delta-frequency modulations) play a crucial role even in non-sensory regions for the modulation of faster frequencies guiding attention (Szczepanski et al., 2014). Here we extend the findings from visuospatial attention tasks (Szczepanski et al., 2013) to the audiovisual domain, showing that frontoparietal regions can also flexibly entrain to temporal properties of behaviourally relevant information.

## 5. Conclusion

The present study confirms and extends previous studies showing that attention can act on the frequency and both the phase and power of entrained oscillations. Critically, we show a differential pattern for primary sensory regions and non-sensory attentional systems. While selective attention modulated the strength and timing of stimulus-driven oscillations in a largely modality-specific manner in primary sensory areas, frontoparietal regions (including the DAN) in general flexibly adjusted the frequency of the entrained oscillation to the selected sensory modality. Whereas our study used highly artificial stimulus settings (for a review on similar experiments see Calderone et al. (2014)), it will be interesting in future studies to scrutinize rich conditions with more naturalistic stimuli like communication. Further investigating naturalistic human interaction is needed to understand how attention is extracting relevant information when presented with more sophisticated input.

## 7. Acknowledgements

This work is supported the University of Salzburg, #158/2017 and by the Austrian Science Fund, P31230.

